# In-Chip Volumetric Printing of Collagen-I Scaffolds for Perfusable and Stretchable Mammary Tissue Models

**DOI:** 10.64898/2026.07.06.736675

**Authors:** Amelia Hasenauer, Kristian Ivkovic, Samuel Thalmann, Bin Wang, Marcy Zenobi-Wong

## Abstract

Engineered epithelial models require three-dimensional extracellular matrix environments that support organized cell growth and allow independent access to luminal and basal compartments. However, many organ-on-chip (OoC) fabrication strategies rely on planar geometries, non-native materials, or multi-step assembly workflows that limit architectural complexity and experimental control. Here, we report a direct in-chip volumetric printing strategy for fabricating stretchable and perfusable collagen-I scaffolds inside custom OoC devices. A vitamin C-regulated ruthenium/sodium persulfate photocrosslinking system enabled high-fidelity printing of collagen-I into open-lumen architectures with ductal- and alveolar-inspired features. By generating scaffolds directly within the final culture device, this workflow eliminates post-print transfer and integrates defined collagen architectures with compartmentalized fluidic access and a mechanically actuable chip format. To support chip-based culture, printed collagen constructs were stabilized after fabrication using EDC/NHS chemistry, which limited thermally induced collagen densification, improved shape retention, and maintained scaffold anchorage during perfusion. The chip design provided separate access to the printed lumen and surrounding basal compartment, which enabled compartment-specific fluid handling while preserving scaffold integrity during inflation, stretching, and perfusion of the printed construct. On the collagen-I scaffolds, human milk-derived mammary epithelial cells formed epithelial layers with tight junctions and lactation associated markers. The platform further supported perfusion culture, in situ staining, and whole-chip volumetric imaging. Together, this work establishes direct in-chip collagen-I volumetric printing as a biofabrication strategy for creating perfusable epithelial tissue chips with native matrix architecture and compartmentalized fluidic control.

## Introduction

Epithelia are organized as polarized cell layers that separate a luminal compartment from the surrounding stromal microenvironment while regulating exchange across the epithelial barrier.[1] The mammary gland is a dynamic example of this organization, with a branched ductal–alveolar network composed of luminal and basal mammary epithelial cells.[2] During lactation, luminal cells in the alveoli synthesize and secrete milk components, while surrounding contractile basal cells promote milk ejection into the connected ductal system.[2,3] Milk production is therefore coupled to dynamic events on both sides of the epithelial barrier: secretions accumulate and are removed during feeding cycles, while basal-facing hormonal cues change in response to infant suckling and lactation stage.[3] Although mammary organoids and biofabricated scaffolds can reproduce selected aspects of lactogenic differentiation, they often lack independent control over basal-facing stimulation and luminal sampling.[4,5] This creates a need for engineered platforms that combine relevant tissue geometries with matrix-inspired microenvironments and controlled perfusion.[5,6]

Organ-on-chip (OoC) platforms are microengineered culture systems that recreate selected tissue features under perfusion.[7] They enable compartmentalized stimulation and flow, as shown in perfusable human mini-colon models where engineered geometry and asymmetric biochemical stimulation improved epithelial organization, long-term maintenance, and functional readouts.[8] To date, many OoC platforms rely on soft lithography and polydimethylsiloxane (PDMS) molding as accessible and low-cost routes for rapid prototyping.[9] However, they often restrict perfusable structures to planar channel layouts and limit the ability to reproduce mammary-relevant features such as curved lumens and branched ducts embedded within native extracellular matrix (ECM).[5,10] Additionally, many conventional chip workflows require multi-step assembly and post-fabrication handling, such as bonding, alignment, tubing connection, and sequential matrix loading, which can increase the risk of leakage, contamination, and device-to-device variability.[11] These limitations motivate direct in-chip printing of native matrix materials to integrate mammary-relevant geometry, matrix cues, and compartmentalized flow within a single device.

Three-dimensional (3D) bioprinting is increasingly combined with microfluidic systems to create biomimetic hydrogel architectures directly within perfusable devices.[12] Volumetric printing (VP) is well suited for this purpose because it uses computed light projections to generate a 3D dose distribution within a photosensitive volume, to enable the rapid fabrication of complex hydrogel structures under cell-compatible conditions.[13,14] This is particularly promising for chip-integrated scaffold fabrication, where structures must be patterned inside confined device chambers.[15,16]

Previous work has demonstrated the potential of VP for mammary tissue engineering by generating ductal and alveolar scaffolds from decellularized extracellular matrix, which supported human milk-derived mammary epithelial cells (milk MECs) and lactation-associated readouts.[5] Building on this biological framework, collagen-I offers a more defined material platform for engineering mammary-relevant scaffolds with improved compositional control. Collagen-I is a major component of the mammary extracellular matrix (ECM) and provides native cell-adhesive and structural cues, making it an attractive material for defined, cell-instructive 3D architectures.[17,18] However, many printable collagen systems rely on chemical modification to improve printability and shape stability, which can alter native biochemical and fibrillar features.[19,20] We recently addressed this limitation by introducing vitamin C as a biocompatible redox regulator in a ruthenium-mediated photo-crosslinking system, enabling high-fidelity VP of collagen-I without prior chemical modification (C-Redox system).[21,22] These advances create an opportunity to extend collagen-I printing from free-standing scaffolds toward chip-integrated, perfusable matrix architectures.[16] Although existing mammary models can reproduce aspects of lactogenic behavior and OoC systems can provide perfusion, there remains a need for strategies that fabricate native ECM and mammary-like architectures directly inside perfusable chips while preserving separate luminal and basal-facing access.

Here, we introduce an in-chip VP strategy for fabricating 3D collagen-I scaffolds directly within the cylindrical chamber of custom organ-on-chip devices (VP-OoC) (Figure 1). Unlike conventional on-chip approaches, which often pattern cells or matrices on planar chip surfaces, this workflow generates defined 3D collagen architectures inside the final culture device, eliminating post-print transfer. [21] By adapting vitamin C-regulated collagen printing to confined chip chambers, the platform combines native matrix cues with tissue-inspired geometry and compartmentalized luminal and basal-side perfusion. To support stable in-chip culture, scaffold stabilization strategies were developed to modulate collagen densification while maintaining a mechanically robust and cell-compatible scaffold architecture. Using milk MECs, we demonstrate that in situ printed collagen-I architectures can support perfusable epithelial culture within a VP-OoC device.[5] More broadly, this work establishes a generalizable strategy for direct in-chip fabrication of perfusable collagen scaffolds.

**Figure 1:**
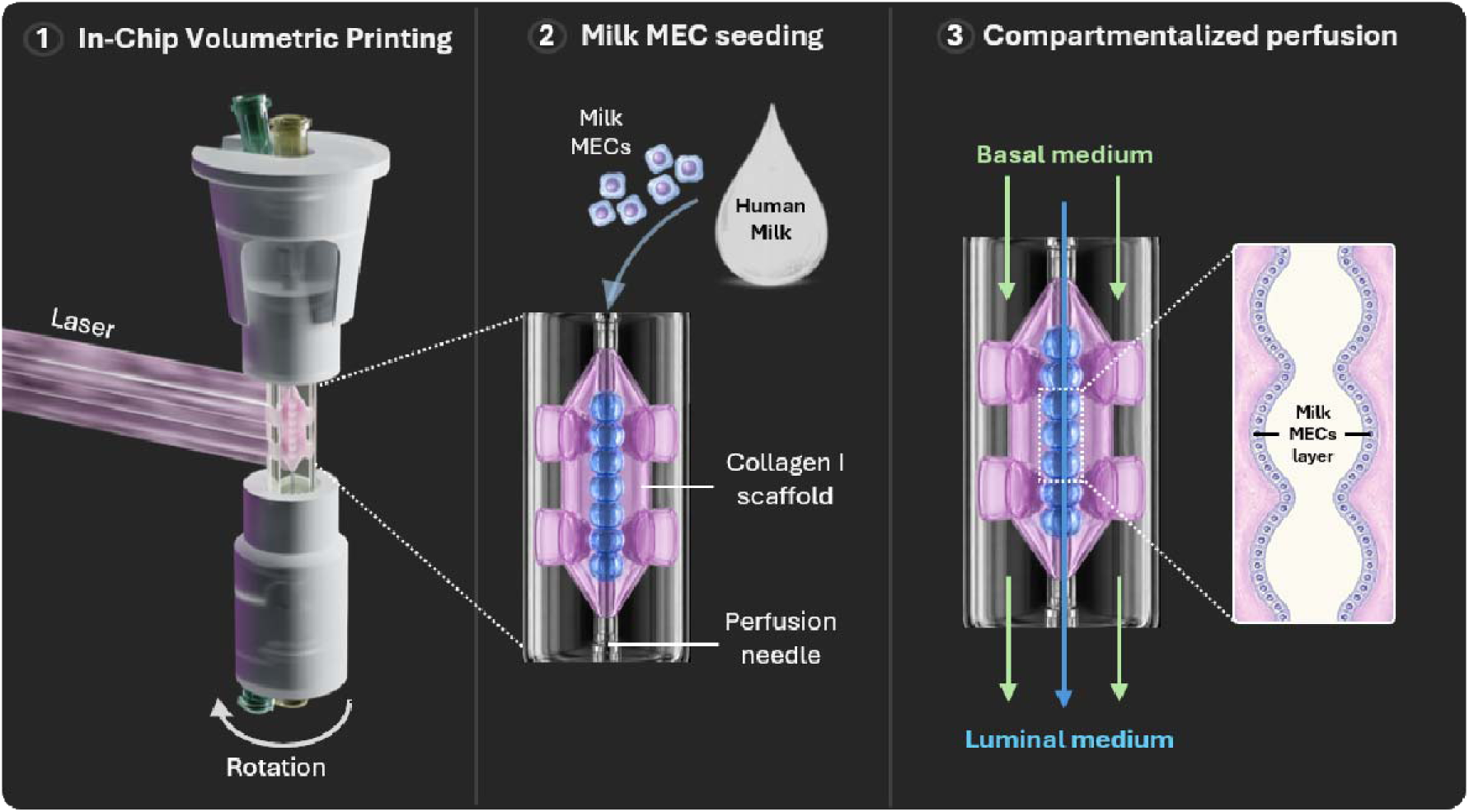
In-chip volumetric printing of collagen-I for compartmentalized mammary epithelial culture. **1)** Chemically unmodified collagen-I is volumetrically printed directly inside the custom-made chip to form a scaffold within the final culture device. **2)** Primary human milk-derived mammary epithelial cells (milk MECs) are seeded onto the collagen-I scaffold through perfusion needles. **3)** The platform enables compartmentalized perfusion, with basal medium delivered to the stromal compartment and luminal medium perfused through the central channel.

## Results

### Evaluation of collagen-I formulations for chip-compatible VP

The C-Redox system introduces a vitamin C-regulated dose threshold for Ru/SPS-mediated collagen-I crosslinking, enabling spatially confined VP (Figure 2A).[21] Based on this chemistry, collagen-I concentration and light dose were evaluated to define a working range for direct in-chip printing of lumenized collagen architectures with sufficient structural stability for post-processing and perfusion.

**Figure 2:**
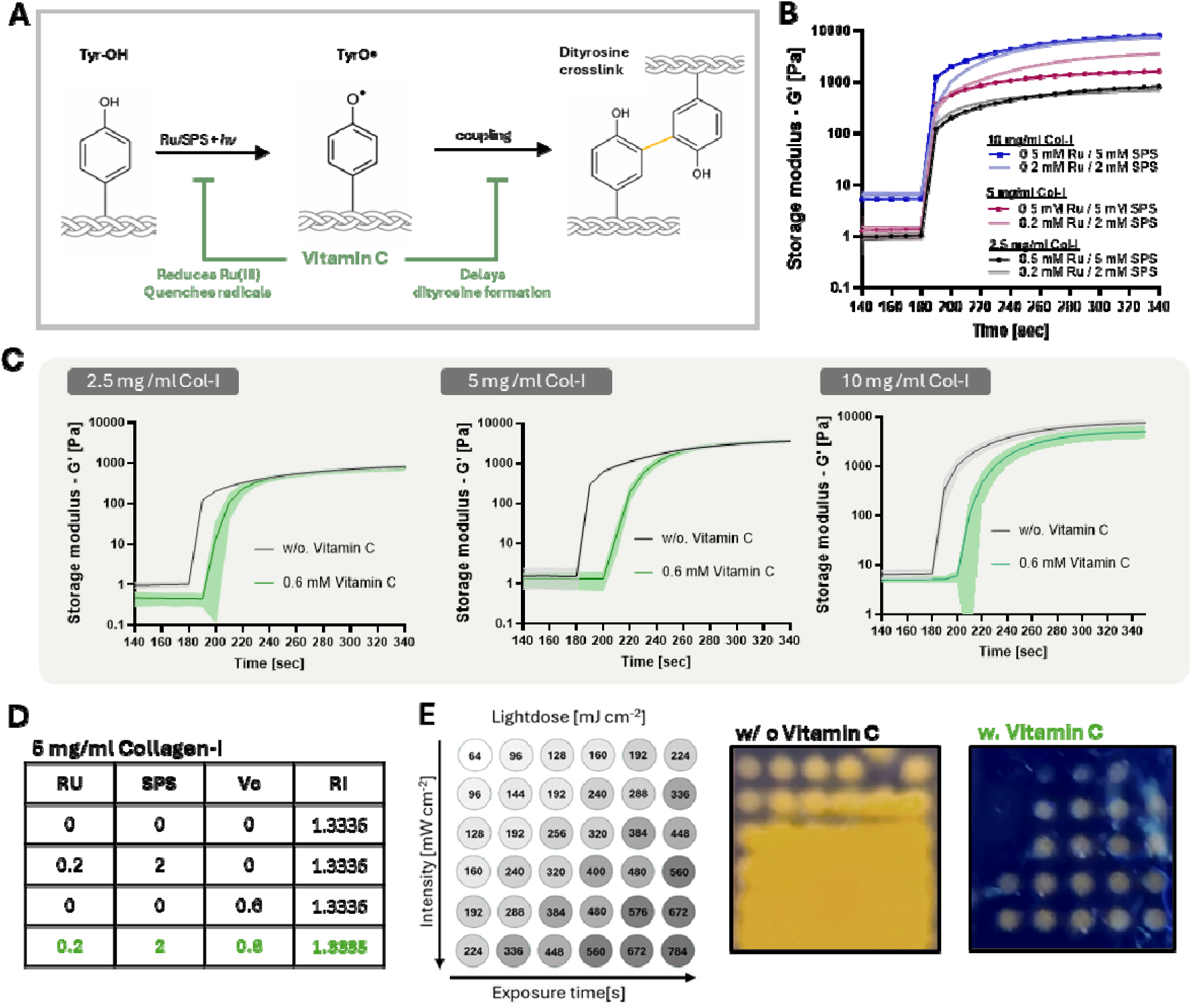
Collagen-I formulation screening for chip-compatible VP. **A)** Schematic of the C-Redox system for Ru/SPS-mediated collagen-I photocrosslinking, in which vitamin C delays radical-driven dityrosine crosslinking. **B)** Photorheological screening of collagen-I crosslinking across collagen-I and Ru/SPS concentrations. **C)** Photorheological evaluation of 2.5, 5, and 10 mg/mL collagen-I with 0.2 mM Ru/2 mM SPS, with or without 0.6 mM vitamin C. **D)** Refractive index measurements of 5mg/mL collagen-I containing Ru/SPS, vitamin C, or the complete C-Redox system. **E)** Light-dose test of 5 mg/mL collagen-I with and without vitamin C across increasing exposure times and light intensities. Without vitamin C, collagen-I showed diffuse gelation, whereas C-Redox formulations produced spatially confined dots. Data are shown as mean ± SD where applicable.

Ruthenium/sodium persulfate (Ru/SPS)-mediated crosslinking was first assessed by photorheology across different collagen-I and photoinitiator concentrations (Figure 2B). Light exposure induced rapid gelation in all formulations, with plateau storage moduli increasing with collagen-I concentration from ∼0.7–0.9 kPa at 2.5 mg/mL to ∼2–3 kPa at 5 mg/mL and ∼5–7 kPa at 10 mg/mL. Increasing Ru/SPS above 0.2 mM Ru and 2 mM SPS did not substantially increase the plateau modulus at any tested collagen concentration, indicating that gel stiffness was mainly governed by collagen content. Therefore, 0.2/2 mM Ru/SPS was retained for subsequent experiments.

Vitamin C regulation was then evaluated at fixed Ru/SPS concentrations in 2.5, 5, and 10 mg/mL collagen-I formulations (Figure 2C). Addition of 0.6 mM vitamin C delayed gelation onset across all collagen concentrations while preserving a concentration-dependent increase in storage modulus. Photorheology with increasing vitamin C concentrations showed a dose-dependent delay in gelation onset, which suggested that gelation timing was more sensitive to redox-regulator concentration than to collagen concentration, whereas final stiffness remained mainly determined by collagen content (**Figure S1**). Based on these results and in line with previous work, 0.6 mM vitamin C was selected because it provided a clear gelation delay while maintaining robust crosslinking.[21]

Based on the photorheology results, 5 and 10 mg/mL collagen-I formulations were selected for further evaluation because their higher storage moduli compared with 2.5 mg/mL indicated greater resistance to deformation, which was expected to improve shape retention of self-supporting, perfusable structures during post-processing and perfusion. Optical compatibility was then assessed by refractive index measurements. The refractive index was higher for 10 mg/mL collagen-I than for 5 mg/mL collagen-I, with values of ∼1.3345–1.3350 and ∼1.3335–1.3340, respectively, and the addition of Ru/SPS, vitamin C, or the complete C-Redox system caused only minor changes in refractive index (Figure 2D and **Figure S2A**).

A light-dose test was performed to identify suitable C-Redox conditions for VP (Figure 2E). Without vitamin C, 5 and 10 mg/mL collagen-I showed diffuse gelation throughout the exposed region, consistent with insufficient suppression of background crosslinking. In contrast, the C-Redox formulation reduced background gelation and produced localized patterns at sufficient light doses. For subsequent in-chip VP, 5 mg/mL collagen I with 0.2/2 mM Ru/SPS was selected as the primary working formulation because it supported reliable pattern formation while reducing collagen use compared with 10 mg/mL (**Figure S2 B-C**). Light doses of 600–750 mJ/cm² were then used for printing.

### Volumetric printing of chemically unmodified collagen-I scaffolds with open lumen architectures

In VP, computed light projections are delivered into a rotating resin volume, forming 3D structures by volumetric dose accumulation (Figure 3A). Printing experiments focused on adapting C-Redox collagen-I to chip-relevant open-lumen architectures. Using 5 mg/mL collagen-I, 0.2 mM Ru, 2 mM SPS, 0.6 mM vitamin C, and a printing dose of 600 mJ/cm² freeform structures were first printed to assess shape fidelity across diverse geometries such as a pipe, three multistar structures, a lumen structure with support legs, and a milk-carton model (Figure 3B). More complex lumen-containing architectures were then printed, such as non-linear and duct-like channels with alveolar-like side features that reproduced relevant 3D curvatures. Light-sheet imaging confirmed preservation of these internal lumen geometries and continuous open channels (Figure 3C). Perfusion of fluorescently labeled gelatin methacryloyl (GelMA) through bifurcated collagen-I scaffolds further confirmed that the branched lumen network was continuous and accessible (**Figure S3A**).

**Figure 3:**
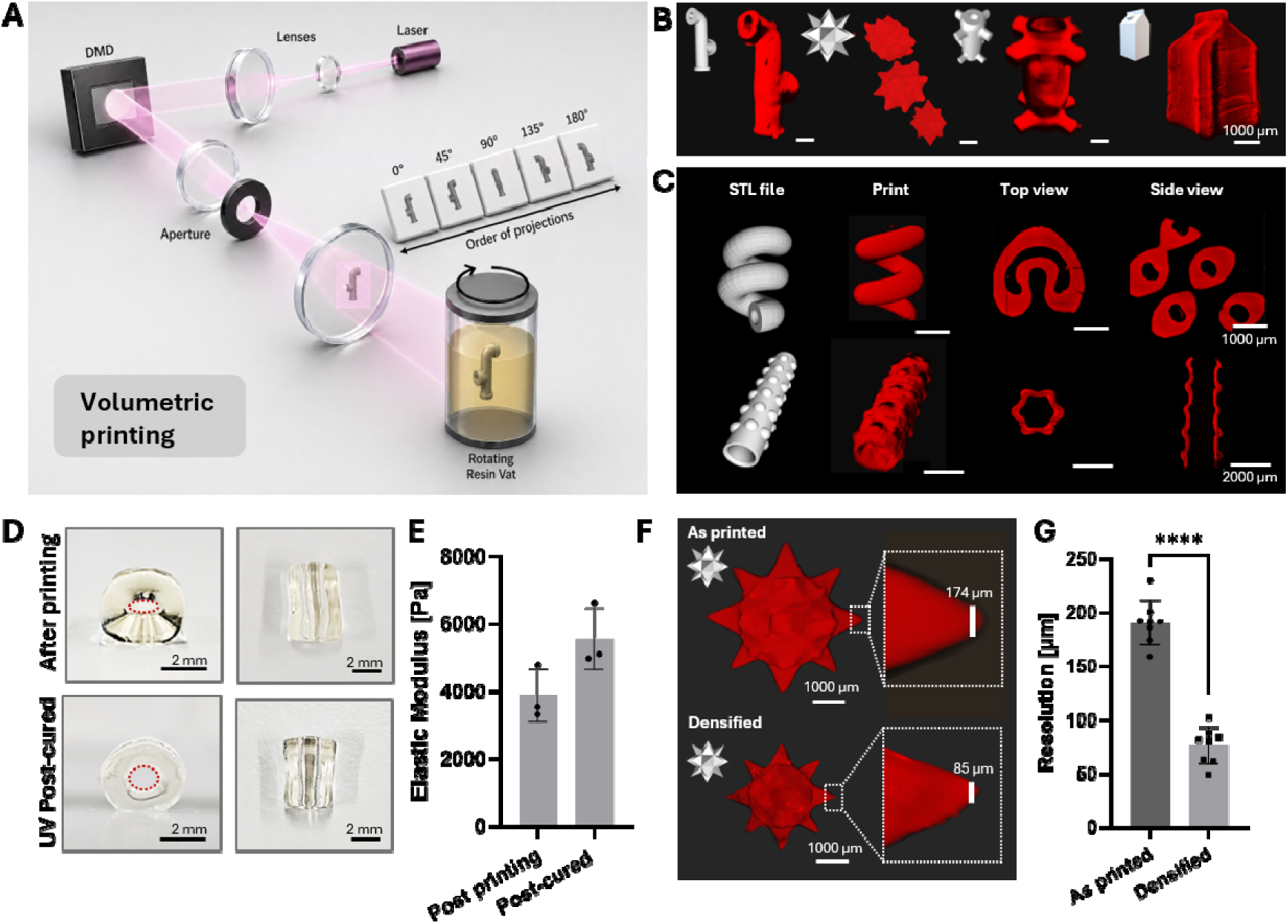
Tomographic volumetric printing of perfusable collagen-I scaffolds. **A)** Schematic of the VP setup, in which computed light projections are delivered into a rotating resin vat to generate three-dimensional collagen-I structures by volumetric dose accumulation. **B)** Representative collagen-I architectures printed with the C-Redox formulation. **C)** Printed collagen-I scaffolds with complex perfusable lumen architectures, such as hollow helices and duct-like structures with alveolar-like features. STL files, whole-print views, top views, and side views are shown where indicated. **D)** Representative collagen-I constructs after printing and after UV post-curing, showing improved retention of macroscopic geometry and open lumen structures. Red dotted lines indicate the lumen outline. Scale bars: 2 mm. **E)** Elastic modulus of collagen-I constructs after printing and after UV post-curing. Data are shown as mean ± SD, n = 3 technical replicates. **F)** Representative fluorescence images of a printed collagen-I resolution structure before and after densification. Densification reduced the representative feature size from 174 µm to 85 µm while preserving overall geometry. Scale bars: 1000 µm. **G)** Quantification of feature size before and after densification, showing a significant reduction in effective feature dimensions after densification. Data are shown as mean ± SD; ****p < 0.0001, n = 8 technical replicates.

Open lumens that maintain their structure facilitate cell seeding as well as handling of the printed construct and chip. Lumen stability was therefore evaluated across different lumen-to-wall thickness ratios (**Figure S3B**). This screening showed that lumen stability depended on scaffold geometry and supported the use of an approximately 1:1 lumen-to-wall thickness ratio for subsequent chip experiments, as this geometry better maintained open channels during handling. UV post-curing was then introduced to further improve lumen shape retention before seeding and perfusion (Figure 3D and **Figure S3B**). Mechanical characterization showed an increase in elastic modulus after post-curing, from 3–4 kPa to 5–6 kPa (Figure 3E). Although this increase was not statistically significant, it was consistent with the qualitative improvement in lumen shape retention observed after post-curing (**Figure 3D–E**).

An additional dose comparison was performed to assess the upper end of the selected printing window. Collagen-I scaffolds printed at 750 mJ/cm² remained printable but showed larger feature sizes and reduced resolution compared with 600 mJ/cm² but produced scaffolds with a higher elastic modulus after printing (**Figure S4**). Both doses underwent similar densification during post printing incubation at 37°C. This shrinkage increased the feature resolution from <200 µm to <100 µm while preserving overall scaffold geometry (**Figure 3F–G**). Together, these results define chip-compatible printing and post-processing conditions for open-lumen collagen-I scaffolds.

### VP-compatible chip design for in-situ scaffold fabrication and compartmentalized perfusion

To translate free-standing collagen-I architectures into a perfusable culture platform, a custom VP-compatible OoC device was developed. The chip was designed to support sterile culture, provide independent fluidic access to the printed lumen and outer basal compartment, permit sample recovery after culture, and remain compatible with commercial and open-source volumetric printing systems through a top-loading format that allows insertion and removal from above. The device consists of top and bottom caps, a rubber lid, a mixing plug, a round glass printing chamber, and a needle assembly that provides access to the printed scaffold (**Figure 4A–B** and Figure **S5**). The dimensions of the chip are compatible with the Tomolite volumetric printing systems (**Figure S6**).

**Figure 4:**
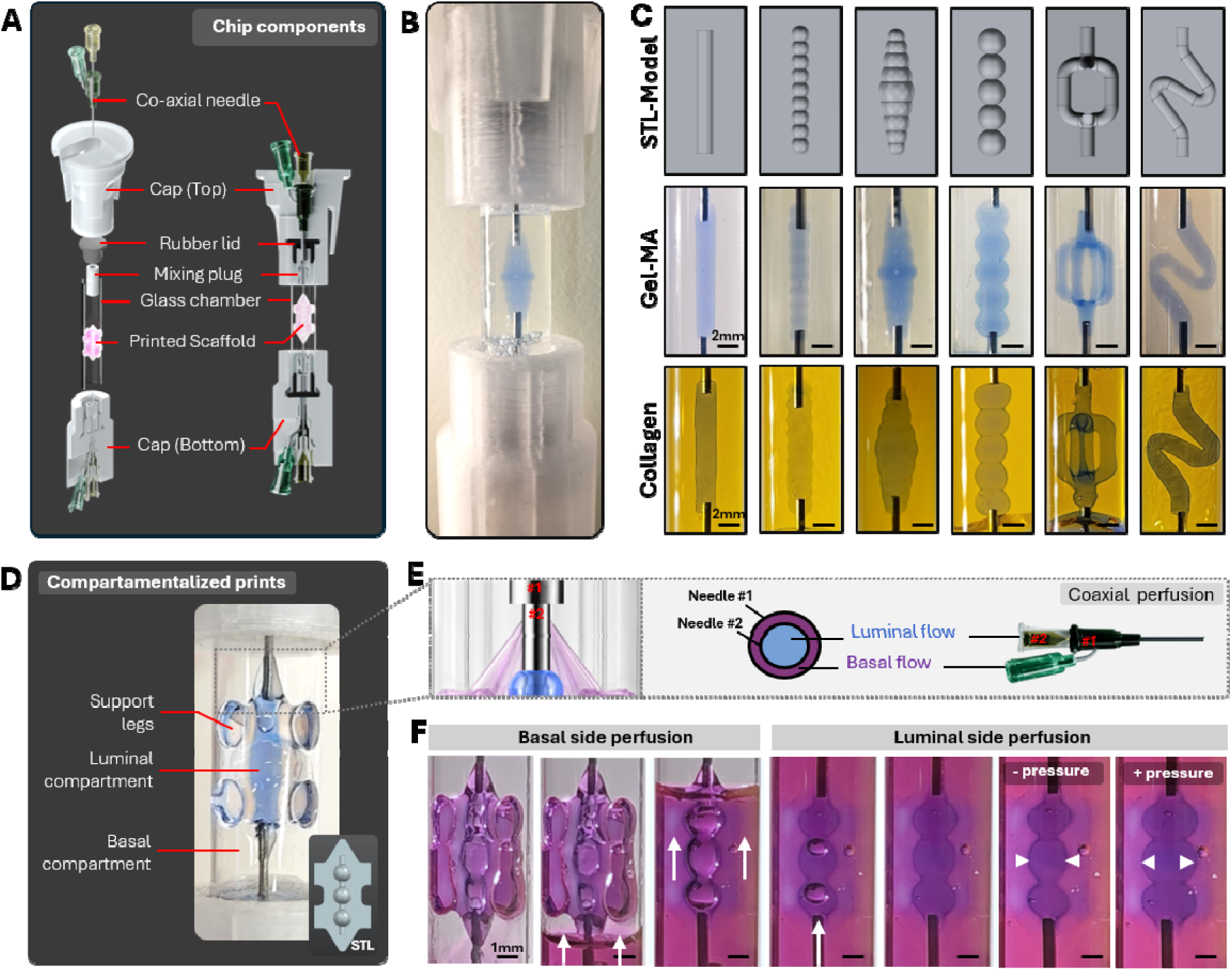
Design and validation of a VP-OoC platform for compartmentalized perfusion of printed collagen-I scaffolds. **A)** Exploded and assembled schematic of the custom microfluidic chip components, including the glass chamber, top and bottom caps, rubber lid, mixing plug, co-axial needle, and printed collagen scaffold. The design enables scaffold fabrication within a confined chamber while maintaining access to luminal and surrounding compartments. **B)** Representative image of the assembled chip chamber containing a printed collagen-I scaffold around the central needle. **C)** Design and in-chip fabrication of collagen-I scaffolds with different lumen geometries. STL models, printed constructs, and corresponding brightfield images show successful generation of straight, helical, branched, bead-like, and duct-like scaffold architectures inside the chip chamber. Scale bars: 2 mm. **D)** Compartmentalized collagen-I scaffold design with support legs, a central luminal compartment, and a surrounding basal/stromal-facing compartment. The inset shows the corresponding STL model. **E)** Schematic of compartment-specific perfusion using the co-axial needle configuration, enabling separate access to the luminal flow path and the basal/stromal-facing compartment. **F)** Basal-side perfusion filled the outer space around the scaffold (treated with EDC/NHS), while luminal-side perfusion filled the internal row of alveolar-like lumens. White arrows indicate perfusate entry and filling; white arrowheads indicate lumen boundaries during negative- and positive-pressure testing.

In-chip printing was first tested with 5% GelMA to confirm that VP could be performed inside the confined glass chamber before translating the workflow to collagen-I. A cylindrical GelMA construct with perfusable internal channels was printed around the needle assembly in the assembled chip (Figure 4B), followed by straight, alveolar-like, branched, and curved lumen designs in both GelMA and collagen-I (Figure 4C).

To move from internal channel printing toward a compartmentalized scaffold, a collagen-I scaffold with a more complex outer geometry was developed and tested with 5% GelMA (**Figure S7A**). Support legs were incorporated to anchor the printed scaffold to the glass chamber walls while maintaining an open space around the construct to create an outer basal perfusion compartment between the scaffold and the chamber wall (Figure 4D). Separate access to the internal lumen and outer basal compartment was then achieved using a coaxial needle system (Figure 4E). The needle assembly provided luminal access to the printed internal channel, while the surrounding flow path enabled perfusion of the outer basal compartment (**Figure S5C–D).**

Compartment-specific perfusion was evaluated by manually introducing colored solution through the separate flow paths of the coaxial needle system. Solution delivered through the basal path filled the outer space surrounding the printed scaffold, whereas luminal perfusion filled the internal row of alveolar-like lumens (Figure 4F and **Figure S7B-C**). In both cases, the dye remained confined to the intended region, which confirmed separate basal and luminal perfusion within the chip-integrated construct. The scaffold maintained its position and overall geometry over the 10 repeated filling and emptying cycles tested, demonstrating stable anchorage during cyclic fluid loading. Additional negative- and positive-pressure tests showed reversible deflation and inflation of the luminal compartment, respectively, with inward and outward displacement of the printed scaffold walls. This pressure-driven deformation was repeatable and occurred without structural failure, which indicate that the chip-integrated collagen construct retained both mechanical compliance and perfusable compartmentalization (Figure 4F, **Videos S1 and S2**). Together, these results support in-chip fabrication of collagen-I scaffolds with compartment-specific perfusion and stability during repeated fluid handling.

### EDC/NHS crosslinking stabilizes VP-printed collagen-I constructs in-chip

Collagen-I densification presents a specific challenge for in-chip scaffold fabrication (Figure 5A). Although densification can improve feature resolution in free-standing VP-printed collagen-I constructs, as shown in **Figure 3F–G**, contraction inside the VP-OoC can compromise scaffold anchorage, lumen alignment, and perfusion compatibility. This is particularly relevant because the construct is constrained by fixed needles and the glass chamber wall. As an initial stabilization strategy, disk-like anchoring features were added to retain collagen-I constructs on the needle assembly during densification. Despite improved construct attachment, shrinkage still distorted the internal lumen geometry (**Figure S8**), indicating that mechanical anchoring alone was insufficient to preserve perfusable scaffold architecture.

**Figure 5:**
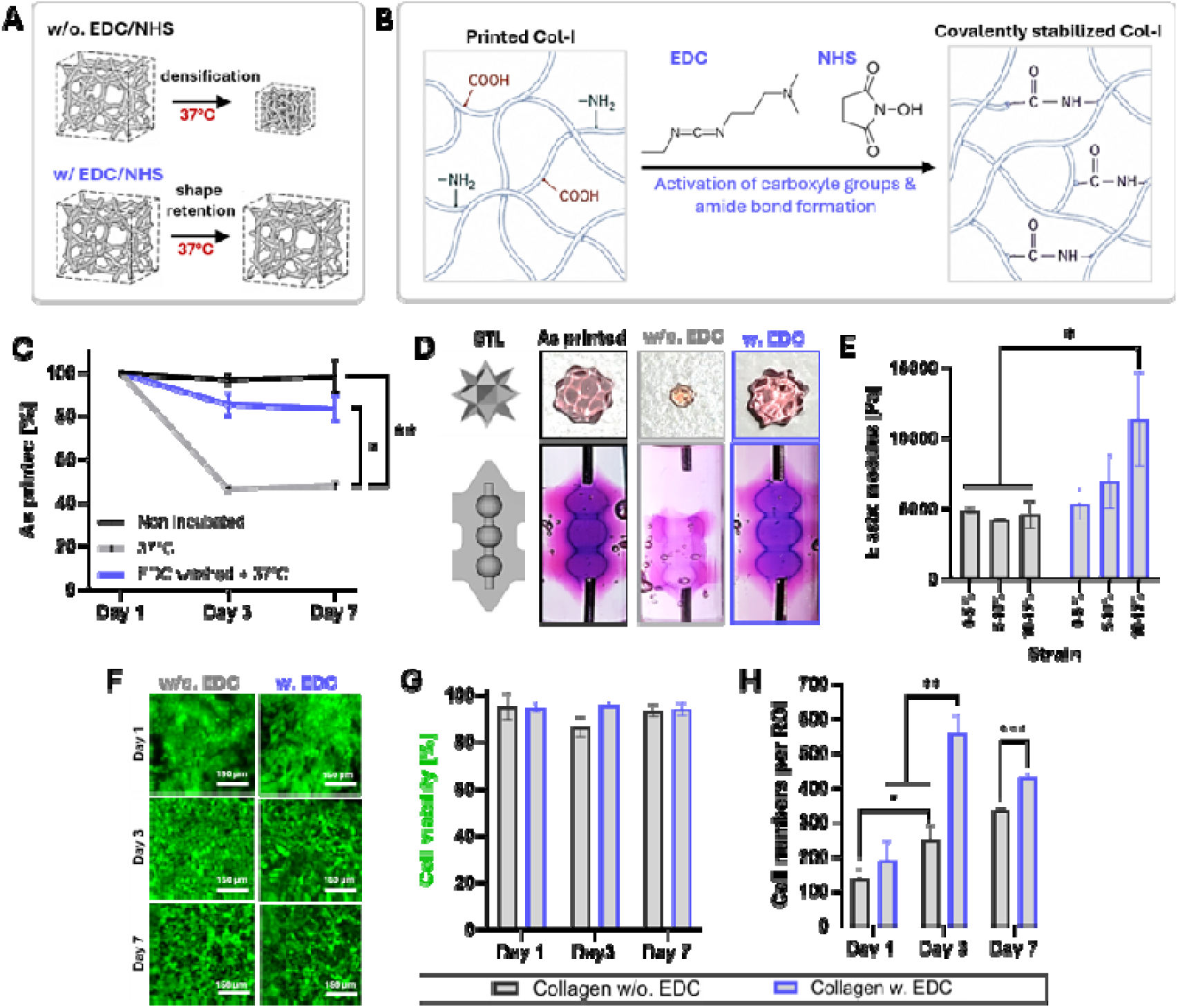
EDC/NHS crosslinking inhibits VP-printed collagen-I densification in-chip and preserves mammary epithelial cell compatibility. **A)** Schematic showing collagen densification at 37°C without EDC/NHS and improved shape retention after crosslinking. **B)** Schematic of EDC/NHS-mediated covalent stabilization of VP-printed collagen-I through carboxyl activation and amide bond formation. **C)** Shape-retention quantification over 7 days for non-incubated, 37°C-incubated, and EDC/NHS-treated and incubated collagen constructs. **D)** Representative images of star-shaped and in-chip collagen constructs showing isotropic densification without EDC/NHS and improved shape retention and needle anchorage after EDC/NHS treatment. **E)** Elastic modulus of untreated and EDC/NHS-treated collagen constructs across increasing strain ranges. **F)** Representative live/dead images of mammary epithelial cells cultured on untreated and EDC/NHS-treated collagen constructs for 1, 3, and 7 days. Scale bars: 150 µm. **G)** Quantification of cell viability over 7 days. **H)** Quantification of cell number per region of interest. Data are shown as mean ± SD; statistical significance is indicated in the figure.

Chemical stabilization of the collagen network was pursued using EDC/NHS chemistry.[23] EDC/NHS activates carboxyl groups on collagen and promotes amide bond formation with available amine groups, which results in additional covalent crosslinks within the network (Figure 5B). Shape-retention analysis of star-shaped constructs showed that after 7 days, non-incubated controls maintained ∼100% of their printed size, whereas untreated constructs incubated at 37 °C retained only ∼50%. EDC/NHS flushing before incubation improved shape retention to ∼85% of the printed geometry (Figure 5C and **Figure S9A-C**).

The EDC/NHS stabilization strategy was evaluated in the assembled VP-OoC format. Untreated constructs showed pronounced shrinkage after incubation, resulting in detachment from the fixed needles and chamber wall. In contrast, EDC/NHS-treated constructs remained anchored to the needle assembly and glass chamber, preserved their alignment, and appeared visually comparable to non-incubated controls (Figure 5D). Mechanical testing further showed that EDC/NHS stabilization increased scaffold stiffness. Untreated constructs exhibited elastic moduli of approximately 4–5 kPa across the tested strain range, whereas EDC/NHS-treated scaffolds showed higher elastic moduli, increasing from ∼5 kPa at 0–5% strain to ∼11 kPa at 10–15% strain (Figure 5E).

VP-printed C-Redox collagen-I scaffolds were then evaluated for mammary epithelial cell compatibility, and the effect of EDC/NHS stabilization. MCF10A cells, a non-tumorigenic human mammary epithelial cell line, was seeded onto VP-printed collagen-I constructs. Live/dead staining showed cell attachment in both untreated and 15 min EDC/NHS-treated constructs over 7 days, with high viability of >85% across conditions (**Figure 5F–G**). Cell numbers increased over time in both conditions and were higher on EDC/NHS-treated constructs than on untreated collagen by day 7 (Figure 5H). A longer 60 min EDC/NHS treatment was additionally tested and remained compatible with MCF10A attachment and viability over the 7-day culture period (**Figure S9D**). Together, these results show that VP-printed C-Redox collagen-I scaffolds support mammary epithelial cell culture and that EDC/NHS stabilization limits densification while preserving cell compatibility and in-chip anchorage.

### Perfusion-compatible collagen-I chips support primary human mammary epithelial cell culture

With scaffold stabilization and cytocompatibility established, the platform was advanced from material validation to mammary epithelial tissue modeling. VP-printed collagen-I scaffolds with a ductal–alveolar-inspired geometry were first evaluated under static culture conditions, independent of the chip, to assess mammary epithelial cell organization before implementation in perfusion-compatible devices. Both MCF10A cells and human milk-derived mammary epithelial cells were seeded directly onto densified and EDC/NHS-treated collagen-I scaffolds. F-actin staining showed cell coverage across the scaffold surface in both scaffold conditions and with both mammary cell types (Figure 6A). Cross-sectional views further confirmed expression of mammary epithelial lineage markers, such as cytokeratin 8 (CK8) for luminal epithelial cells and cytokeratin 14 (CK14) marking basal/myoepithelial cells along the collagen lumen. Both densified and EDC/NHS-treated scaffolds remained stable with cells attached throughout the 7-day culture period (Figure 6A and **Figure S10**).

**Figure 6:**
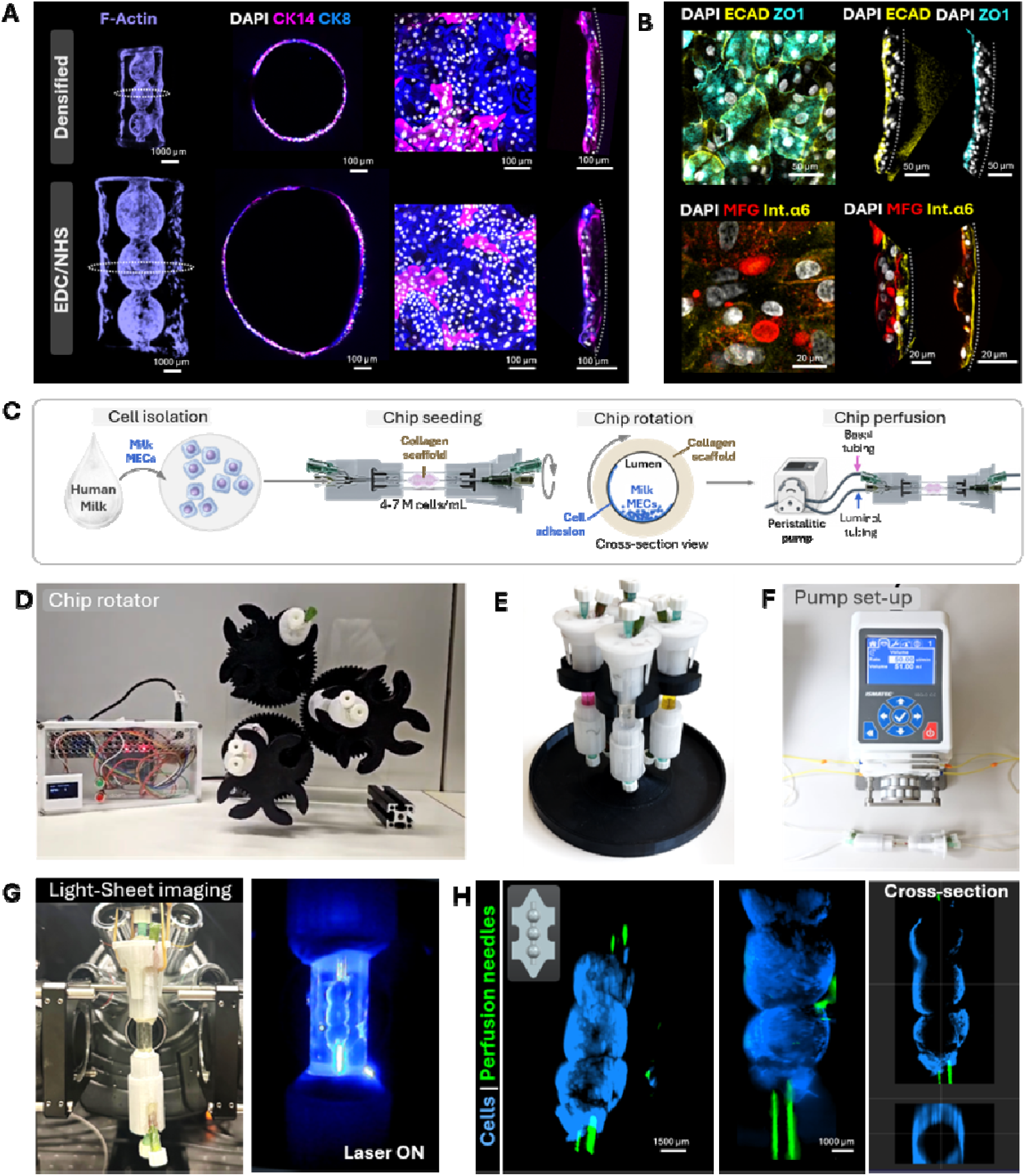
VP-printed collagen-I scaffolds support mammary epithelial cell organization and perfusion-compatible chip culture. **A)** Representative F-actin staining and cross-sectional CK8/CK14 immunostaining of mammary epithelial cells on densified and EDC/NHS-stabilized collagen-I scaffolds. **B)** Immunostaining of EDC/NHS-stabilized scaffolds for E-cadherin, ZO-1, MFG, and integrin _α_6/CD49f. **C)** Workflow for human milk-derived mammary epithelial cell isolation, chip seeding, rotation-assisted attachment, and perfusion culture. **D)** Custom chip rotator for dynamic seeding. **E)** Chip holder for upright culture and parallel device handling. **F)** Peristaltic pump setup for lumen perfusion. **G)** Light-sheet microscopy setup for whole-chip imaging after perfusion culture. H) Three-dimensional reconstructions and cross-sectional views showing mammary epithelial cells around the printed lumen and perfusion needles within the chip. Scale bars are indicated in the panels.

Further immunostaining was performed on EDC/NHS-stabilized collagen scaffolds used for subsequent chip integration. Epithelial cadherin (E-cadherin), an adherens junction protein, and zonula occludens-1 (ZO-1), a tight junction-associated protein, localized along cell–cell interfaces. Milk fat globule (MFG), a marker associated with mammary secretory/luminal differentiation, was observed toward the lumen-facing region, while integrin-α6 (CD49f) was enriched along the matrix-facing side, consistent with basal attachment to the collagen scaffold (Figure 6B).

After confirming mammary epithelial marker expression on VP-printed collagen-I scaffolds under static conditions, the workflow was translated into the chip format. Human milk MECs were seeded into the printed chip lumen, followed by chip rotation to promote circumferential cell attachment before perfusion culture (Figure 6C). A custom chip rotator enabled dynamic seeding by distributing mammary epithelial cells around the printed collagen lumen (**Figure 6C,D**).[24] After an initial attachment period, the chips were transferred to a holder for parallel handling of multiple devices (Figure 6E). Perfusion was then established through the lumen using a peristaltic pump at 50 µL/min for 4 days, supporting cell retention and culture within the collagen-I chip under flow (Figure 6F).

Following perfusion culture, the chips were stained in situ by perfusing the F-actin staining solution through the inner channel. The chip was mounted in a light-sheet microscope to enable whole-chip volumetric imaging without disrupting the printed collagen construct (Figure 6G). This enabled three-dimensional visualization of mammary epithelial cells within the perfused chip and showed their spatial organization relative to the perfusion needles (Figure 6H). Cross-sectional views further confirmed cell localization around the printed lumen after perfusion (Figure 6H). Together, these results demonstrate that VP-printed collagen-I scaffolds support mammary epithelial cell attachment and organization under static conditions and can be integrated into perfusion-compatible chip cultures suitable for whole-chip volumetric imaging.

## Discussion

Engineered epithelial models are increasingly shaped by advances in biofabricated scaffolds and OoC technologies.[7] Yet it remains difficult to combine native matrix composition, predefined three-dimensional architecture, and compartmentalized perfusion within a single platform. This challenge is particularly relevant for secretory epithelia such as the mammary gland, where luminal and basal-facing cues regulate tissue organization and function.[2,25] Here, we establish direct in-chip VP of collagen-I as a strategy to bridge this gap. By adapting C-Redox collagen-I printing to confined chip chambers, we generated mammary-inspired scaffolds directly inside the final culture device, eliminating post-print transfer while preserving separate luminal and basal access. These printed collagen architectures supported primary human milk MECs attachment and epithelial marker expression in a perfusable VP-OoC platform.

Within collagen biofabrication, methods such as FRESH have shown that native collagen can be printed into complex, perfusable structures, but typically require support baths, nozzle-based deposition, and post-print handling.[16,26] Light-based printing offers rapid, contact-free fabrication, but often relies on modified collagen, gelatin-derived materials, or synthetic photoresins rather than collagen.[27,28] Here, collagen-I is combined with VP directly inside a perfusable chip. This preserves native matrix composition while using VP to define scaffold geometry and fluidic access within the final culture device.

This direct in-chip workflow also creates opportunities for more adaptive fabrication strategies. In the current setup, scaffold alignment with the coaxial needles and perfusion paths still depends on manual positioning, which may affect lumen placement and connection quality. Recent advances in context-aware and overprinting-based VP suggest practical ways to make this alignment more automated and reproducible. Generative, adaptive, context-aware volumetric bioprinting (GRACE) demonstrates how imaging-guided model adaptation can adjust print geometry to the real print environment, while displacement- and refraction-corrected tomographic volumetric additive manufacturing (Dr.TVAM) shows that projection patterns can be optimized around pre-existing or optically complex features, including chip interfaces.[29,30] In a related preprint, Rizzo et al. applied TVAM directly inside preassembled microfluidic chips, using Dr.TVAM-guided projection planning to account for the optical constraints of square chip geometries.[31] For the present platform, these advances could improve alignment with needle interfaces, support more complex inlet and outlet designs, and potentially enable collagen architectures to be printed around pre-positioned organoids or epithelial aggregates within perfusable chips.

For chip-integrated collagen scaffolds, shape stability is essential because deformation can disrupt perfusion and fluidic connections. In conventional microfluidic collagen models, gel deformation is often managed through device-level strategies such as confinement, phaseguides, surface adhesion, or hydrogel anchoring.[32,33] These approaches are effective for localizing bulk hydrogels, but they are less suited to the present challenge, where a predefined printed lumen must retain its shape and remain connected to fixed needle interfaces throughout washing, incubation, and cell seeding.[32] Hydrogel anchoring was therefore considered, but network-level stabilization was required to limit deformation of the printed collagen architecture itself. EDC/NHS stabilization addressed this by reinforcing the collagen network and improving shape retention within the chip. Carbodiimide crosslinking may also alter matrix mechanics and cell–matrix interactions.[23,34] The subsequent attachment of primary human milk MECs indicates that EDC/NHS-treated scaffolds can support post-seeding culture of a comparatively sensitive primary epithelial cell source.[5] However, the effects of this stabilization step on collagen bioactivity and potential cell-laden printing remain to be further assessed.

The compartmentalized design is particularly relevant because mammary secretion is regulated across two sides of a polarized epithelial barrier. In the lactating gland, mammary epithelial cells secrete proteins, ions, and lipid droplets into the alveolar lumen, while lactogenic hormones and stromal-derived cues act from the basal side to regulate differentiation and secretory activity.[35–37] A single perfused channel could support luminal clearance, but it would not independently control the basal-side environment. The separated luminal and basal compartments of the VP-OoC platform therefore enable a more physiologically relevant experimental design, in which basal stimulation can be varied while secreted products are collected from the lumen over time. In addition to sample collection, luminal flow may help remove locally accumulated secretory material and provide mechanical cues to the epithelial layer, although the effects of shear on mammary epithelial polarization and secretion remain to be tested directly.[38,39] The next biological step is therefore to use the platform to couple basal lactogenic stimulation with luminal flow and collection, moving from static epithelial attachment toward functional mammary secretion studies.

In summary, this work advances VP-OoC engineering by applying direct in-chip printing to pristine collagen-I scaffolds for mammary epithelial culture. C-Redox collagen printing was combined with EDC/NHS stabilization to preserve native collagen composition while maintaining scaffold geometry within fixed microfluidic interfaces. The final platform integrates mammary-inspired collagen architecture with separate luminal and basal compartments, to enable epithelial cell culture in a perfusable chip format. This approach moves mammary epithelial modeling beyond static scaffold culture toward systems where basal stimulation and luminal collection can be studied together and provides a route to more functional mammary secretion models.

## Materials and Methods

### Photorheology

Rheological measurements were performed using an Anton Paar MCR302e rheometer with a 20 mm parallel-plate geometry and a 0.2 mm gap. Samples were irradiated through a 405 nm light filter at an irradiance of 4.02 mW/cm² and maintained at 25 °C using a LAUDA Alpha RA 8 cooling thermostat. Resins were freshly prepared in a dark room before each measurement, and 76 µL was loaded onto the stage. A damp tissue was placed around the sample to minimize dehydration. Measurements were conducted at 2% shear strain and 1 Hz for 20 min, with light exposure initiated after 3 min. Data were collected every 10 s, with three replicates per formulation. Replicates were plotted as mean ± error. The inhibition period and crosslinking slope were extracted from the curves, while the plateau storage modulus was calculated by averaging values over the final 5 min.

### Compressive Testing

The compressive modulus of printed constructs was measured using a TA.XTplus texture analyzer. Disks were printed with 5 mm in diameter and 2 mm in height for measurements. Measurements were performed using a 500 g load cell and a 15 mm cylindrical probe. Samples were centered on the stage, preloaded with 0.2 g to ensure contact, compressed to 15% strain at 0.1 mm/s, and decompressed at the same rate. Moduli were determined for the 5–10% and 10–15% strain ranges. Dissipated energy was calculated as the difference between the integrals of the loading and unloading curves.

### Refractive Index

The refractive index was measured with a Kern Optics ABB refractometer. 100 µL of photoresin was placed on the lower prism and covered with the upper prism. The refractive index was determined visually by centering the light–dark boundary in the eyepiece and reading the corresponding value from the calibrated scale.

### Resin Preparation

Stock solutions were prepared in deionized water, including 50 mM ruthenium (Advanced BioMatrix, 5248-1KIT), 500 mM sodium persulfate (SPS; Advanced BioMatrix, 5248-1KIT), and 60 mM L-ascorbic acid (vitamin C; Sigma-Aldrich, 95210). Ruthenium stocks were protected from light with aluminum foil. Stocks were stored at 4 °C for <2 weeks. One day before printing, purified type I bovine collagen (Symatese, CBPE2US) was dissolved in dHLO at 10 mg/mL and solubilized overnight at 4 °C. Resins were prepared in a dark room on ice. The collagen stock was diluted to 5 mg/mL with dHLO, followed by addition of 1% (v/v) vitamin C, 0.4% (v/v) ruthenium, and 0.4% (v/v) SPS, resulting in final concentrations of 0.6 mM vitamin C, 0.2 mM ruthenium, and 2 mM SPS. The resin was mixed by pipetting for 1 min, centrifuged for 5 min at 1000 rcf to remove bubbles.

### Dose Tests

400 µL of resin was loaded into rectangular cuvettes with a 2 mm path length. Dose test parameters are listed in TableS1 printed grids were generated using light doses ranging from 64 to 784 mJ.cm^-^², with the two grid dimensions corresponding to light intensity and exposure time. After printing, dose tests were washed with cold PBS and a rhodamine B solution and imaged using a Samsung Galaxy S23+ phone camera under bright-field or UV illumination inside a photo box.

### Volumetric Printing

Volumetric printing was performed using a Readily3D Tomolite v2 open-format printer operated with Apparite software (ver. 0d96231). Prints were performed in 10 mm diameter glass vials or directly within the chip. Unless otherwise stated, prints were produced using a light dose of 600 mJ.cm^-^² and an average intensity of 15.0 mW.cm^-^². Printing time, number of turns, rotation speed, and projection rate were set using the Apparite “optimal setting” function. Resin volume depended on construct size, ranging from 0.6 mL for smaller structures, such as multistars, to 1.5–1.8 mL for larger perfusable channels. For in-chip printing, the entire glass chamber was filled with 2 mL resin.

### Post-printing Processing

After printing, vials were removed from the printer and flushed with cold PBS to remove partially crosslinked resin and, when necessary, detach the construct from the vial. Constructs were transferred to 6-well plates containing fresh PBS and washed repeatedly to remove uncrosslinked material and ensure channel patency. Samples were then immersed in fresh PBS and post-cured in a UV box at a light dose of 10 J/cm².

### Secondary Crosslinking

Secondary crosslinking was performed using freshly prepared 10 mM EDC hydrochloride (Fluorochem, 024810) and 5 mM NHS (Sigma-Aldrich, 56485) in dHLO. Samples were immersed in the EDC/NHS solution for 15–30 min with stirring or repeated perfusion, then thoroughly washed with PBS.

### Chip assembly

Top and bottom caps were printed on a Prusa MK4 using a 0.4 mm nozzle, 0.15 mm layer height, 15% infill, and supports. The mixing plug and needle cup were printed using a 0.25 mm nozzle, 0.12 mm layer height, 15% infill, supports, and a brim. Needle assemblies were prepared by cutting 1-inch 14G Metcal dosing needles to 12 mm, smoothing the cut edges with 180-grit sandpaper, and drilling a centered lateral hole with a 1.5 mm drill bit. A 0.5-inch 14G 45° angled Metcal dosing needle was inserted into the lateral hole while maintaining an unobstructed axial lumen. Each needle assembly was bonded into the cross-shaped recess of the chip cap using epoxy. After initial curing, epoxy was applied to the junction between the straight and angled needles. A 21G × 50 mm BD Microlance needle was inserted through the straight dosing needle during curing to maintain alignment, while the angled needle was held vertically. Silicone rubber plugs were centrally punctured. The glass tube was wetted and inserted into the chip using a slow screwing motion to minimize fracture.

### Needle polydopamine coating

To further improve the adhesion, Stainless-steel needles were roughened and functionalized with a polydopamine coating before printing. Needles were cleaned by immersion in 70% ethanol for 10–15 min, rinsed thoroughly with deionized water, and dried under sterile conditions. A dopamine coating solution was freshly prepared by dissolving dopamine hydrochloride at 2 mg/mL in 10 mM Tris-HCl buffer, pH 8.5. The needles were fully immersed in the dopamine solution and incubated for 2–18 h at room temperature under gentle agitation.

### In-Chip Printing

For in-chip printing, the two bottom nozzles were sealed with Luer-lock screw caps. The glass chamber was inserted into the lower chip cap, filled with resin, and closed with the upper cap. One upper outer nozzle was left open during assembly to allow pressure equalization and escape of excess air or resin. The assembled chip was inserted directly into the Readily3D printer without a collet and secured with the clamping nut. After printing, the channels were washed with cold PBS using syringes connected to the respective inlet and outlet nozzles. In-chip constructs were then UV post-cured and treated with EDC/NHS as described above.

### 3D Design

Scaffolds constructs and FDM-printed accessories were designed in Siemens NX CAD software (version 2406 3000). Chip components were designed and modified in Autodesk Fusion (version 2605.0.97).

### Fused Deposition Modeling (FDM) Printing

FDM printing was performed using an Original Prusa MK4 with standard settings and enabled supports. Larger components were printed using the “Original Prusa MK4 Input Shaper 0.4 nozzle” preset with “0.15 mm STRUCTURAL” settings, while smaller components, such as mixing plugs, were printed using the “Original Prusa MK4 Input Shaper 0.25 nozzle” preset with “0.12 mm STRUCTURAL” settings. Chip components were printed from PolyLite PETG Clear, and other accessories from PolyLite PETG Black.

### Light-Sheet Imaging

Printed constructs were imaged with an axially scanned light-sheet microscope (MesoSPIM, V4). Constructs were immersed overnight with 5 mg/mL Rhodamine B in PBS; densified constructs were immersed in Rhodamine MA during densification. Samples were transferred to 4 mm glass cuvettes containing Milli-Q water, mounted on a custom 3D-printed holder, and submerged in a Milli-Q-filled quartz chamber on the microscope stand. Imaging was performed with an Olympus MVX-10 macro-zoom system and MVPLAPO1x 2× air objective with adjustable zoom. Electrically tunable lens voltage was adjusted for each run, and step sizes ranged from 5 to 50 µm. Images were post-processed and visualized in Imaris Viewer x64 9.9.1

### Immunofluorescent staining and confocal imaging

Printed constructs seeded with cells were fixed with 4% paraformaldehyde (PFA) 8 h. After three PBS washes, samples were permeabilized and blocked in PBS containing 5% bovine serum albumin (BSA; MilliporeSigma) and 0.2% Triton X-100 (Sigma-Aldrich, T8787) at room temperature. Primary antibodies (Table S2) were applied overnight at 4°C in BSA-PBS. Samples were then washed with PBS and incubated for 2 h at room temperature with secondary antibodies (Table S3) and 4′,6-diamidino-2-phenylindole (DAPI) diluted in BSA-PBS. After final PBS washes, samples were imaged using an Olympus FV4000 confocal microscope.

### Resolution Analysis

Printing resolution of complex constructs was assessed using bright-field imaging on an EVOS M5000 microscope. The pointed tips of printed star constructs were imaged, and positive resolution was quantified as the distance between the two points where the printed tip deviated from a linear edge and began to round.

### Densification Analysis

Densification was assessed using printed star-shaped constructs. Unless used as controls, constructs were incubated at 37 °C in cell culture medium for at least 16 h. Samples were imaged using a Leica Wild Heerbrugg M650 microscope, and inner diameters were measured in the Leica software from eight angular positions per construct to account for measurement variability. Mean diameters were calculated in Excel (Microsoft 365 MSO, Version 2412, Build 16.0.18324.20092, 64-bit).

### Cell culture

MCF10A cells (CRL-10317, ATCC) were expanded in MCF10A medium as previously reported.[40] Cells were cultured at 37 °C with medium changes every 2 days until approximately 85% confluency. Human milk-derived mammary epithelial cells (milk MECs) were obtained from consenting donors in accordance with Swiss Ethics guidelines (Kantonale Ethikkommission, 2022-02012). Cryopreserved human milk-derived mammary epithelial cells (milk MECs), previously isolated from consenting donors under Swiss Ethics approval (Kantonale Ethikkommission, 2022-02012), were thawed and expanded for this study. Cells were cultured in Mammary Epithelial Cell Growth Medium (MECGM; PromoCell, C-21010) supplemented with 10 µM forskolin, 5% human serum, and 1% Pen/Strep at 37 °C and 5% OL. Cells were passaged at 80–90% confluency using TrypLE (Gibco, 12604021) and reseeded at 6000 cells.cm^-^².[5,41]

### Cell Seeding

Disks, perfusable channels, and in-chip perfusable scaffolds were seeded. For flat disks, 40,000 cells in 10 µL were pipetted onto each construct. After 45 min incubation at 37 °C, constructs were washed with media and returned to culture. Perfusable channels were seeded at the same density and incubated for 45 min, with 90° rotation every 10 min to promote uniform channel coverage. In-chip scaffolds were seeded by injecting the same cell suspension through the inner luminal nozzle, followed by incubation for 2.5 h on a rotation device programmed to rotate 90° every 5 min.

### Viability

Cell viability was assessed using Hoechst 33342 (Invitrogen, H3570), Calcein AM (Invitrogen, C3099), and propidium iodide (Fluka, 81845-25MG). Samples were incubated in staining solution for 40 min at 37 °C and washed with PBS. Images were acquired using an EVOS M5000 microscope.

### Quantification and statistical analysis

Experiments were performed with three or more replicates, unless otherwise specified. The sample size (*n*) for each experiment is indicated in the corresponding figure legends. Data analysis was performed using Excel (Microsoft 365 MSO Version 2412, Build 16.0.18324.20092, 64-bit), MATLAB R2018a (version 9.4.0.813654), GraphPad Prism (version 10.4.0 for Windows), and Fiji/ImageJ 1.54f (Java 1.8.0_332). Unless stated otherwise, data are presented as mean ± SD. Statistical significance was assessed using a paired *t*-test, with *P* < 0.05 considered statistically significant. Only statistically significant comparisons are indicated in the figure panels, unless otherwise specified in the figure legends.

## Acknowledgements

We are grateful to the participating mothers for donating milk samples. Thank you to Jakub Janiak for sharing GelMA. We thank Ann-Sophie Frind for advice on EDC/NHS chemistry, Simon Moser for helpful discussions, and all lab members for their valuable input. We also acknowledge ScopeM for support and assistance throughout this work. Light-sheet imaging was performed using equipment maintained by the Center for Microscopy and Image Analysis at the University of Zurich. Funding from the ETH Zürich Foundation (23-1 ETH-012) is gratefully acknowledged.

## Data Availability Statement

All data needed to evaluate the conclusions in the paper are present in the paper and/or the Supplementary Materials. Data also is available in the ETH Zurich Research Collection (https://doi.org/10.3929/ethz-c-000801754) under the terms of the repository’s data-sharing policies.

