## Supplementary Material for "In-Chip Volumetric Printing of Collagen-I Scaffolds for Perfusable and Stretchable Mammary Tissue Models"

Supplementary Figures S1-S10

Tables S1-S3

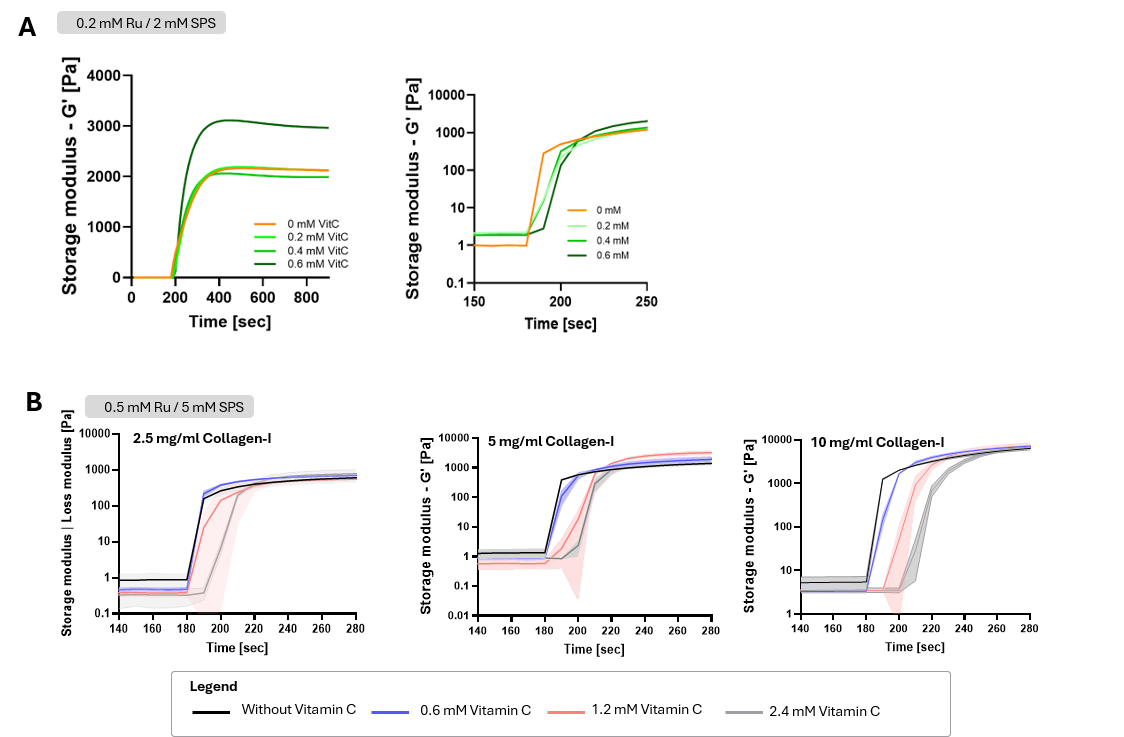

**Figure S1. Vitamin C concentration regulates gelation onset in Ru/SPS-crosslinked collagen-I formulations.** Photorheological analysis of collagen-I formulations containing **A)** 0.2 mM Ru/2 mM SPS or **B)** 0.5 mM Ru/5 mM SPS with increasing vitamin C concentrations. Increasing vitamin C delayed the onset of light-induced gelation under both Ru/SPS conditions, indicating regulation of the gelation threshold. Final storage moduli were mainly dependent on collagen-I concentration. Data are shown as mean ± SD.

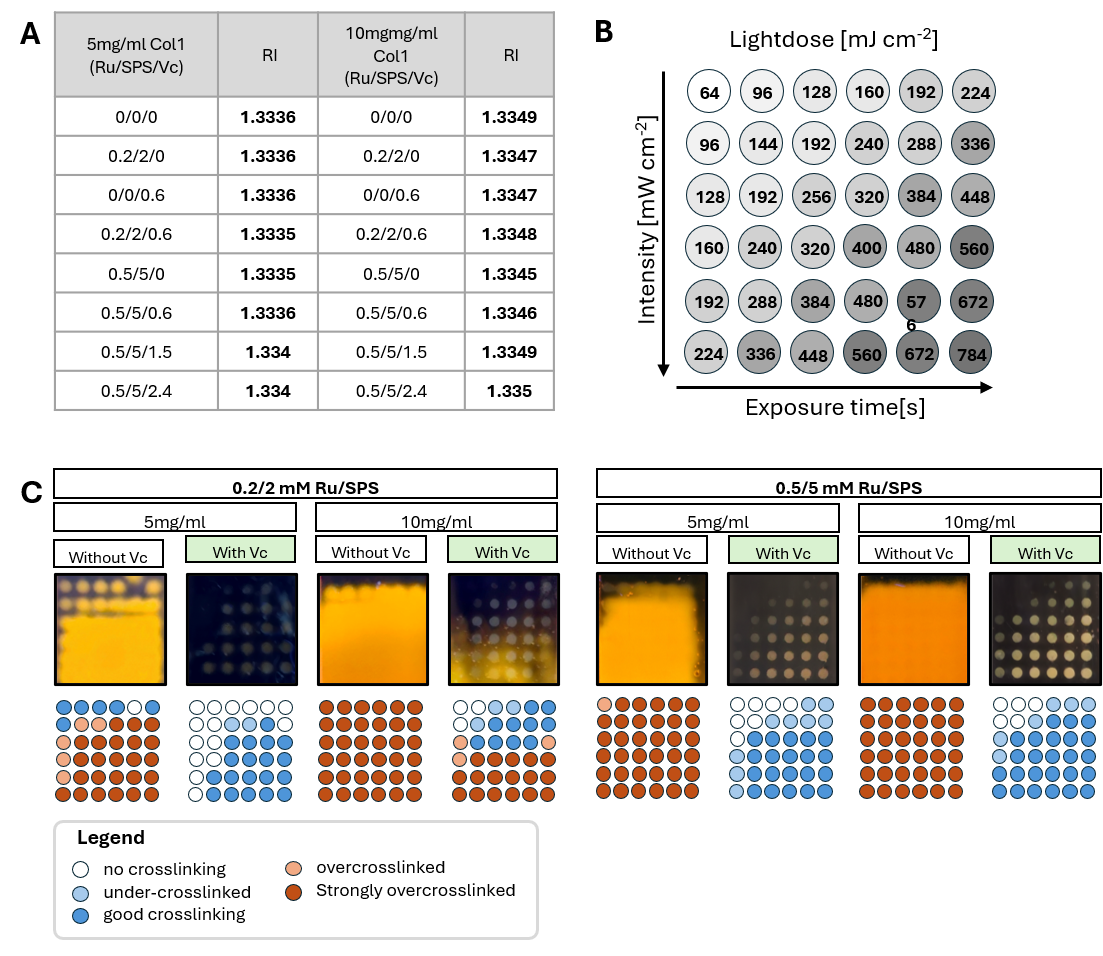

**Figure S2. Refractive index and light-dose screening of collagen-I formulations for VP.** **A)** Refractive index measurements of 5 and 10 mg/mL collagen-I formulations containing different Ru/SPS and vitamin C concentrations. The refractive index was slightly higher for 10 mg/mL collagen-I than for 5 mg/mL collagen-I, while addition of Ru/SPS, vitamin C, or the complete C-Redox system caused only minor changes within each collagen concentration. **B)** Light-dose matrix showing increasing light dose as a function of exposure time and light intensity. **C)** Light-dose screening of collagen-I crosslinking at 0.2/2 mM and 0.5/5 mM Ru/SPS, with or without vitamin C. Vitamin C reduced diffuse background gelation and enabled more spatially confined crosslinking patterns across the tested formulations.

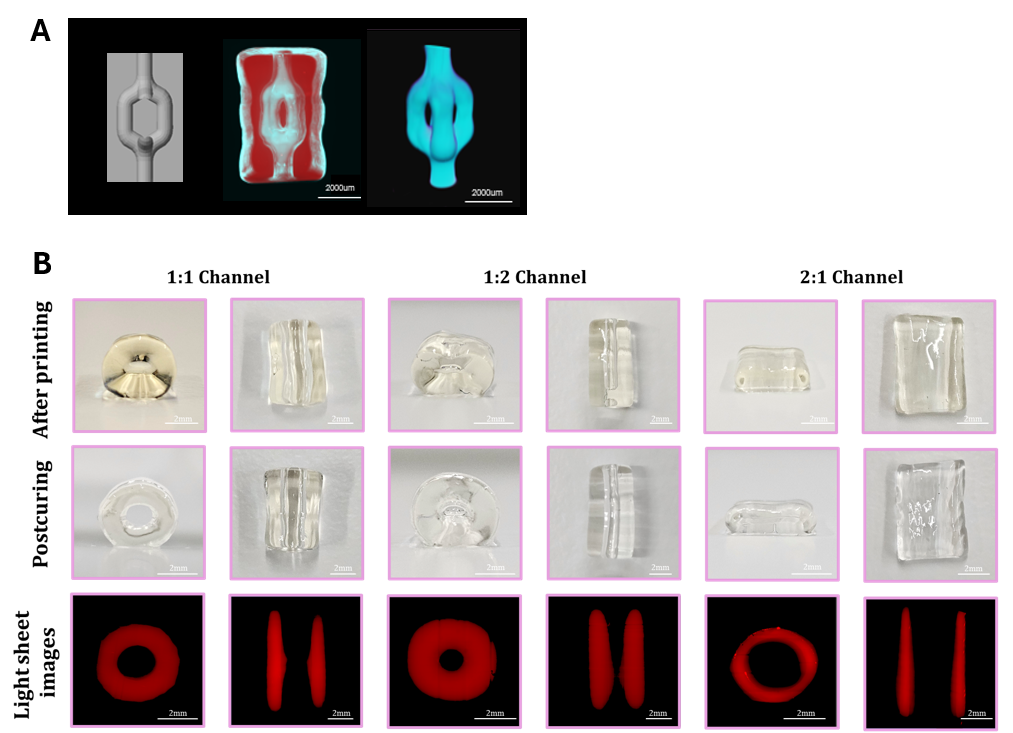

**Figure S3. Perfusion and lumen-stability assessment of printed collagen-I channel architectures.** **A)** Bifurcated lumen scaffold design and perfusion validation. The left panel shows the STL file of the bifurcated channel geometry, the middle panel shows the printed collagen-I scaffold perfused with fluorescently labeled gelatin methacryloyl (GelMA, blue), and the right panel shows the reconstructed perfused channel network. Scale bars: 2000 µm. **B)** Evaluation of collagen-I scaffolds with different lumen-to-wall thickness ratios. Representative photos show constructs after printing and after UV post-curing, while the lower row shows corresponding light-sheet reconstructions of the lumen structures. Scale bars: 2 mm.

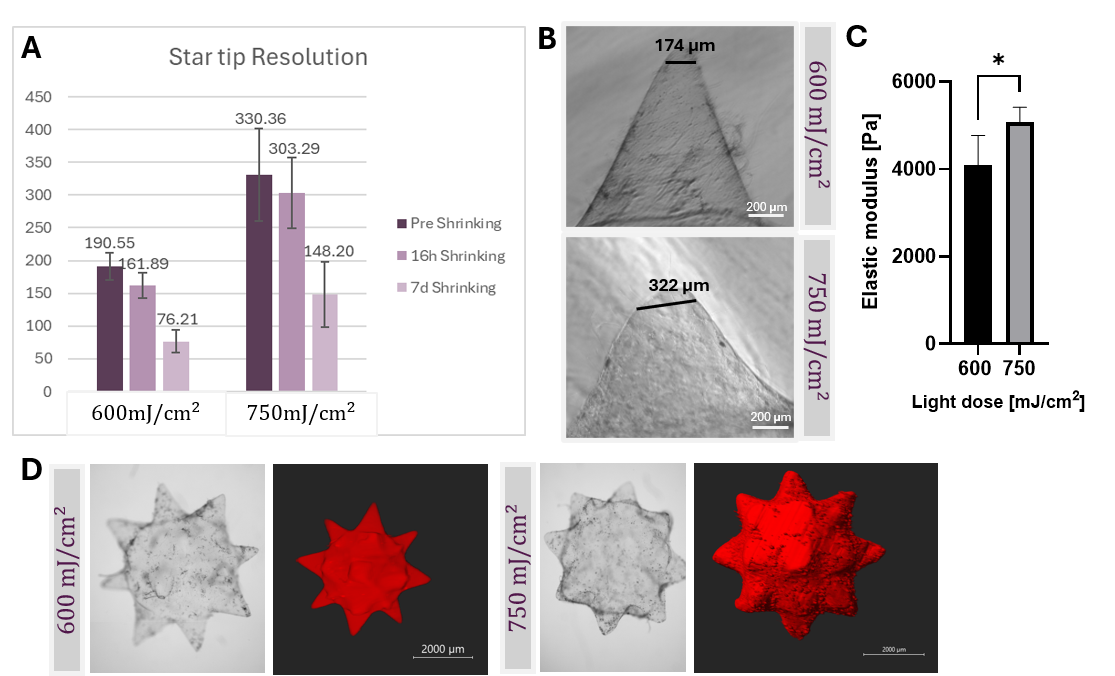

**Figure S4. Effect of printing dose on collagen-I feature resolution and stiffness.** **A)** Quantification of star-tip feature size in collagen-I resolution structures printed at 600 and 750 mJ/cm² before densification, after 16 h densification, and after 7 d densification. **B)** Representative brightfield images of collagen-I resolution structures (star tips) printed at 600 and 750 mJ/cm², with larger feature dimensions at the higher printing dose. Scale bars: 200 µm. **C)** Elastic modulus of collagen-I scaffolds printed at 600 and 750 mJ/cm², with increased stiffness at the higher printing dose. Data are shown as mean ± SD; *p < 0.05. **D)** Overview images of the resulting prints (multistar shaped).

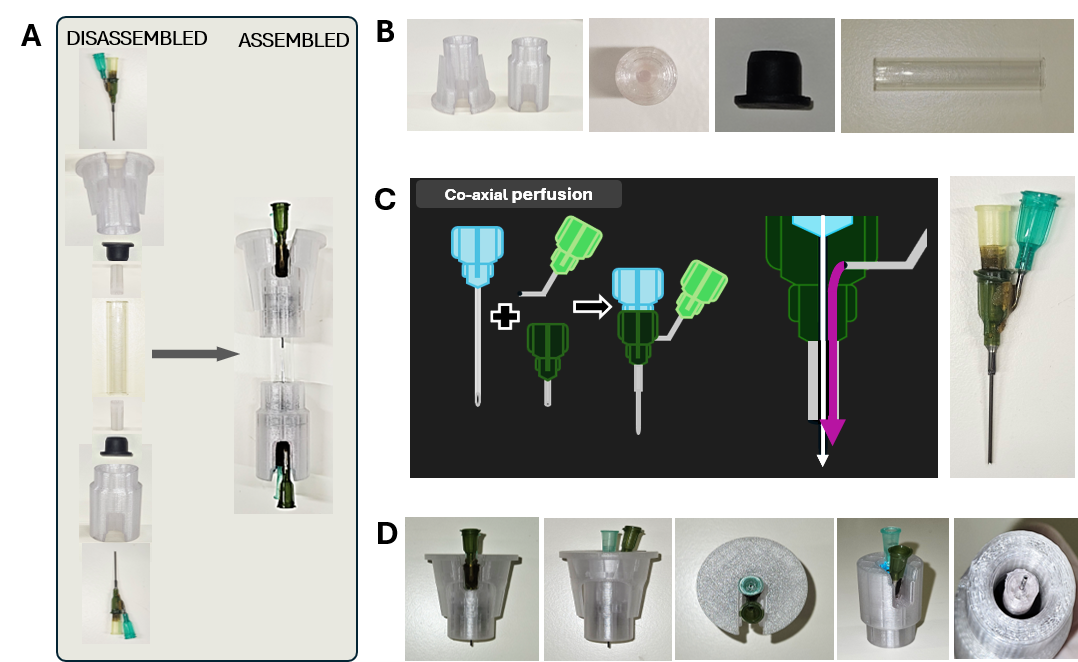

**Figure S5. Assembly and fluidic access design of the VP-compatible organ-on-chip device. A)** Assembled and disassembled views of the VP-compatible chip, showing the main device components and their arrangement around the glass printing chamber. **B)** Close-up images of individual chip components, including the top and bottom caps, rubber lid, mixing plug, glass chamber, and needle assembly. **C)** Schematic and representative image of the coaxial needle setup used to provide separate fluidic access to the printed lumen and outer basal compartment. **D)** Integration of the coaxial needle and mixing plug into the cap system for chip assembly and perfusion access.

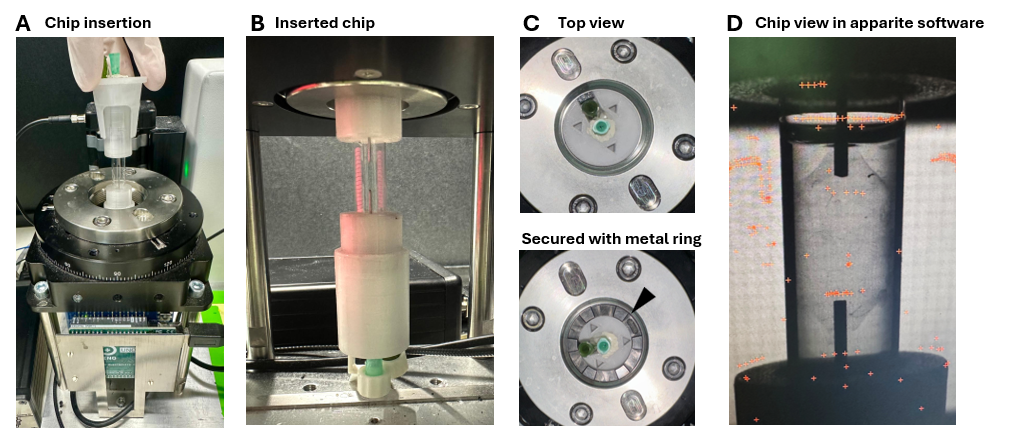
**Figure S6. Top-loading integration of the VP-compatible chip into the Tomolite volumetric printer.** **A)** Representative image of the assembled chip inserted into the Tomolite volumetric printer from above. **B)** Side-view image showing chip alignment within the printing chamber. **C)** Top-view images showing positioning of the chip opening within the printer holder. **D)** Optical visualization of the needle assembly during in-chip printing. The chip was designed for insertion and removal from the top, requiring only vertical access to the printer chamber. This top-loading format enables compatibility with the open-format Tomolite setup shown here and is intended to remain compatible with commercial volumetric printer configurations that provide top access.

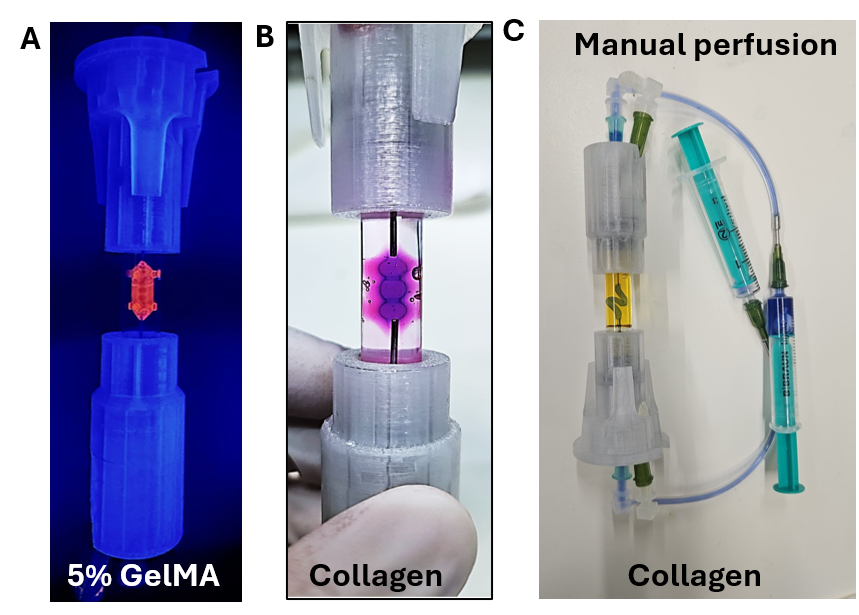

**Figure S7. OoC-VP and manual perfusion setup.** Representative images of the OoC-VP device containing GelMA hydrogel, followed by collagen hydrogel, and the corresponding manual perfusion set-up. The setup illustrates hydrogel integration within the OoC-VP platform and the manual perfusion arrangement used for construct washing before attachment to the peristaltic pump.

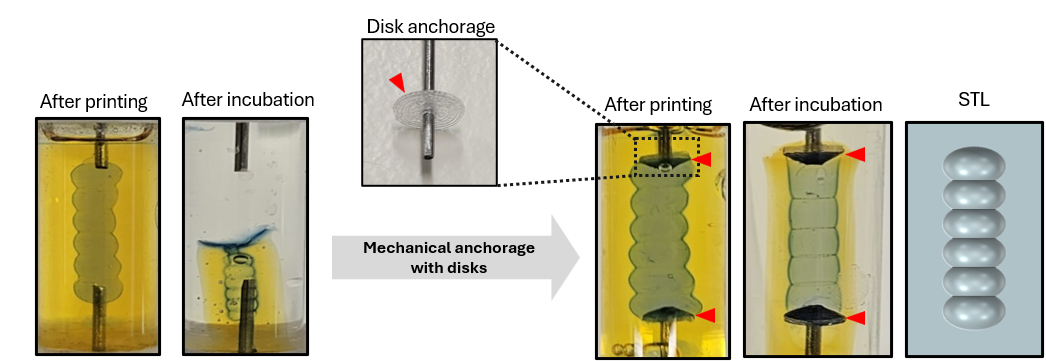

**Figure S8. Mechanical anchoring of densifying collagen-I constructs within the chip.** Representative images showing an initial mechanical anchoring strategy to retain collagen-I constructs during densification. Disk-like anchoring features were printed around the needle assembly to improve construct attachment during shrinkage. Red arrowheads indicate the disk anchorages. Although the anchoring features improved construct retention, densification still distorted the internal lumen geometry, indicating that mechanical anchoring alone was insufficient to preserve the perfusable scaffold architecture.

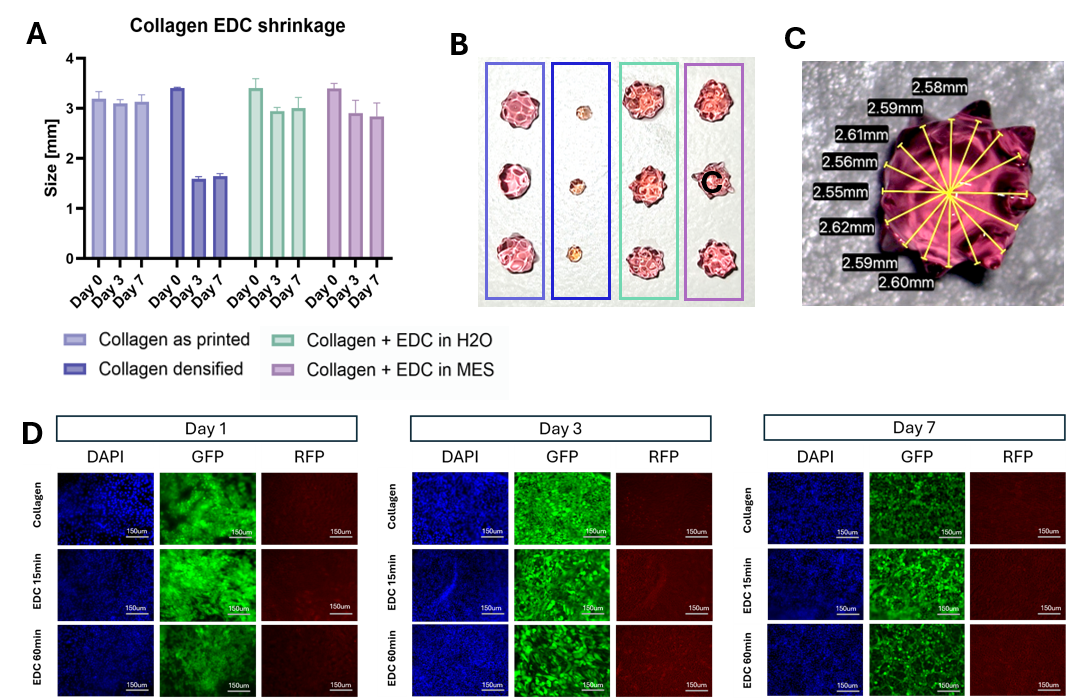

**Figure S9. EDC/NHS stabilization reduces collagen-I scaffold shrinkage and remains compatible with MCF10A culture.** **A)** Quantification of VP-printed collagen-I construct size over 7 days under different stabilization conditions. Untreated collagen constructs underwent pronounced densification during incubation, whereas EDC/NHS treatment in either H₂O or MES buffer improved size retention. Data are shown as mean ± SD. **B)** Representative macroscopic images of collagen-I constructs corresponding to the conditions quantified in A. **C)** Representative radial measurement approach used to quantify construct dimensions and assess size retention across the scaffold geometry. **D)** Cytocompatibility assessment of untreated collagen-I scaffolds and scaffolds treated with EDC/NHS for 15 or 60 min. MCF10A cells were cultured on the constructs for 1, 3, and 7 days and imaged in the DAPI, GFP, and RFP channels. Cells attached to all scaffold conditions and remained predominantly viable over the 7-day culture period. Scale bars: 150 µm.

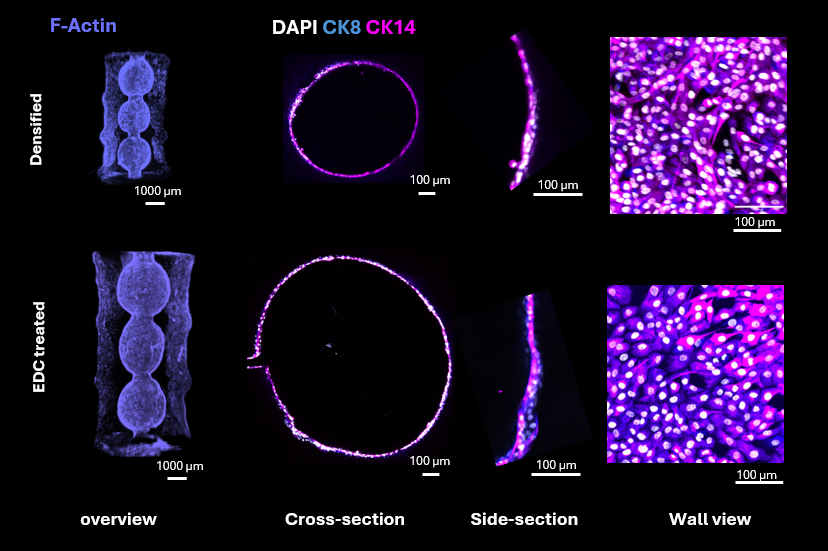

**Figure S10. MCF10A attachment and epithelial marker expression on VP-printed collagen-I scaffolds.** Representative F-actin staining and CK8/CK14 immunostaining of MCF10A cells cultured on densified and EDC/NHS-treated VP-printed collagen-I scaffolds. Overview images show F-actin-labeled cell coverage across the scaffold architecture, while cross-sectional, side-sectional, and wall-view images show the distribution of CK8- and CK14-positive cells along the collagen scaffold wall.

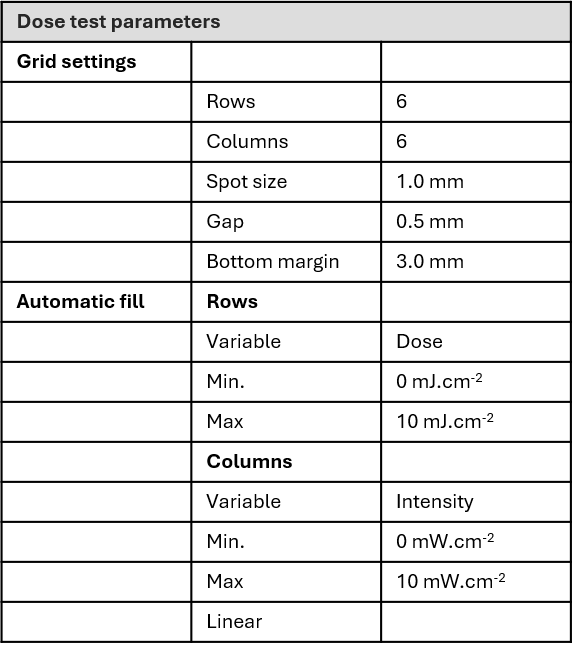

**Table S1.** Dose test parameters.

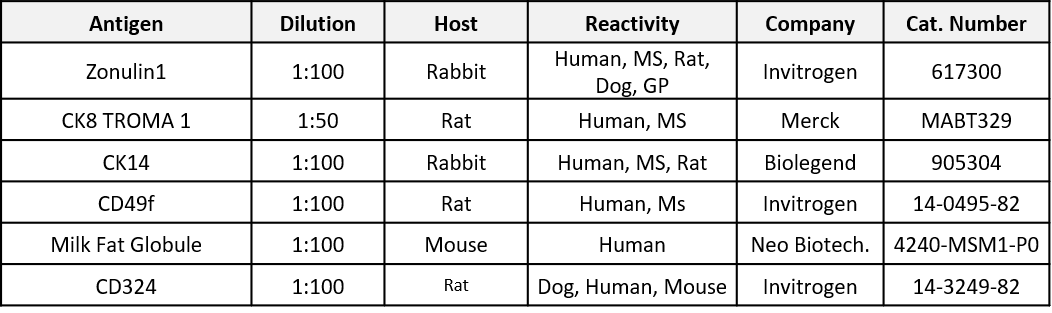

**Table S2.** List of primary antibodies used for immunostaining

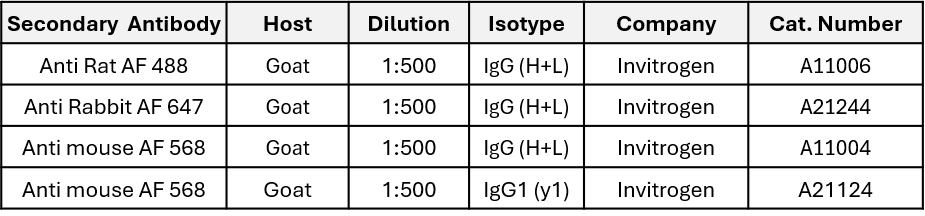

**Table S3.** List of secondary antibodies used for immunostaining
